# Mapping the glial transcriptome in Huntington’s disease using snRNAseq: Selective disruption of glial signatures across brain regions

**DOI:** 10.1101/2022.09.10.507291

**Authors:** Sunniva M. K. Bøstrand, Luise A. Seeker, Nina-Lydia Kazakou, Nadine Bestard-Cuche, Sarah Jäkel, Boyd Kenkhuis, Neil C. Henderson, Susanne T. de Bot, Willeke van Roon-Mom, Josef Priller, Anna Williams

## Abstract

Huntington’s disease (HD) is a severely debilitating, autosomal dominant neurodegenerative disease with a fatal outcome. There is accumulating evidence of a prominent role of glia in the pathology of HD, and we investigated this by conducting single nuclear RNA sequencing (snRNAseq) of human post mortem brain in four differentially affected regions; caudate nucleus, frontal cortex, hippocampus and cerebellum. Across 127,205 nuclei from people with HD, and age/sex matched controls, we found heterogeneity of glia which is altered in HD. We describe prominent changes in the abundance of certain subtypes of astrocytes, microglia, oligodendrocyte precursor cells and oligodendrocytes between HD and control samples, and these differences are widespread across brain regions. Furthermore, we highlight two possible mechanisms that characterise the glial contribution to disease pathology. Firstly, we show that upregulation of molecular chaperones represents a cross-glial signature in HD, which likely reflects an adaptive response to the accumulation of mutant Huntingtin (mHTT). Secondly, we show an oligodendrocyte-specific upregulation of the calmodulin-dependent 3’,5’-cyclic nucleotide phosphodiesterase 1A (*PDE1A*) in HD brain compared to controls, which may cause dysfunction of key cellular functions due to the downregulation of the important second messengers cyclic adenosine monophosphate (cAMP) and cyclic guanosine monophosphate (cGMP). Our results support the hypothesis that glia have an important role in the pathology of HD, and show that all types of glia are affected in the disease. As glia are more tractable to treat than neurons, our findings may be of therapeutic relevance.

## INTRODUCTION

Huntington’s disease (HD) is a rare, autosomal dominant neurodegenerative disorder characterised by choreatic movements, psychiatric and cognitive disturbances^1^. The average age of symptom onset is 40^2^, and the disease typically progresses over a period of up to 20 years before it becomes fatal. HD is caused by a CAG repeat expansion in the first exon of the *Huntingtin (HTT)* gene resulting in expanded polyglutamine (polyQ) repeats in the Huntingtin (HTT) protein. The accumulation of mutant (m) *HTT* RNA and protein causes cellular toxicity by disrupting a number of processes such as proteostasis, transcription and mitochondrial function and the mutant protein forms aggregates in the nucleus and cytoplasm of both neurons and glial cells^3^. The accumulation of *mHTT* is known to result in prominent neurotoxicity and death of the Medium Spiny Neurons (MSNs) of the striatum, disrupting the motor pathways of the basal ganglia and giving rise to the choreatic motor symptoms that are characteristic of HD^4,5^.

While the loss of MSNs is considered the core pathology in HD, all cell types in the body express *HTT*, and there is accumulating evidence that white matter (WM) deficits and changes to glia typically occur in the disease, including astroglia, microglia and oligodendroglia (oligodendrocytes and oligodendrocyte precursor cells (OPCs))^6,7,8,9,10,11^. Global changes to WM microstructure are common in people with HD, as evident both in the neuropathology or on magnetic resonance imaging (MRI), and the latter has been shown to associate with the core symptoms of the disease including motor deficits, cognitive, and psychiatric problems^12,13,14^. WM changes seen on MRI are also observed in premanifest HD, i.e. individuals who are gene expansion-positive but have yet to present with any of the core HD symptoms. Several studies using diffusion tensor imaging (DTI) have demonstrated alterations to WM microstructure at this stage^15,16,17,18^, suggesting that changes to WM represent an early feature of the disease that manifest prior to symptom onset. This in turn hints at the possibility that glial pathology in HD may not simply be secondary to neuronal loss, but may accompany, or even precede, neuronal degeneration. Evidence of oligodendroglial dysfunction in HD, with an onset prior to overt neuronal death has also been shown in several mouse models of HD^19,20^, and different transgenic animal models have been used to investigate the effect of the expanded *mHTT* expression in oligodendroglial populations. Selective expression of *HTT* fragment under the *Plp1* promoter gave rise to a typical motor phenotype similar to the deficits observed in HD, progressive weight loss and premature death^21^. These animals also showed decreased expression of mature oligodendrocyte markers such as myelin basic protein (MBP) and myelin-oligodendrocyte glycoprotein (MOG), as well as thinner myelin sheaths and shortened oligodendrocyte processes.

Ferrari-Bardile et al.^20^ applied the reverse approach, of lowering the expression of *mHTT* under the glial progenitor promoter NG2, thus reducing levels of mHTT in oligodendroglia in a different transgenic mouse model of HD. This was sufficient to rescue myelin deficits in the HD mouse and improve performance in behavioural tests. These studies show that targeting *mHTT* expression in oligodendroglia has a prominent effect on both the cellular and behavioural phenotype in two different mouse models of HD, suggesting that early oligodendrocyte dysfunction in HD has an important contribution to disease pathology that is not secondary to neuronal loss. Furthermore, there was impairment to remyelination after injury as well as developmental myelination in different HD mouse models accompanied by dysregulation of numerous genes downstream of the transcription factor *Tcf7l2*, reflecting a cell-autonomous impairment to oligodendroglia and (re)myelination^22^.

Similar effects of mHTT have also been shown in other glial types. On transplantation of human embryonic stem cell-derived glial progenitor cells (hGPCs) expressing *mHTT* (with the potential to give rise to oligodendrocytes or astrocytes) into myelin deficient mice, these animals produce delayed and disrupted myelin compared to controls^23^. Further, transplantation of hGPCs expressing *mHTT* into healthy mice, led to markedly impaired motor learning and performance as early as 12 weeks^24^. Similarly, transgenic mice with astrocyte-specific expression of *mHTT* also show age-dependent motor function deficits and early death^25^. In a reverse approach, the engraftment of healthy hGPCs (biased towards astroglia by sorting for CD44+) in the R6/2 mouse model of HD, prevented reduction in striatal volume and significantly increased survival, as well as partially rescuing cognitive and motor performance. A similar study to reduce the expression of *mHTT* in mature astrocytes in the BACHD mouse model resulted in a slowing of disease progression, as indicated by slower decline in motor dysfunction and reduced neuropsychiatric-like symptoms as well as rescue of synaptic marker expression and improved neurotransmission in MSNs^26^. In contrast, although reducing *mHTT* levels in brain microglia by 50% rescued the inflammatory phenotype in the BACHD mouse model of HD^27^, this did not prevent the reduction in brain volume nor result in improvement of the HD-like phenotype characterised by motor deficits and anxiety-like behaviour. Conversely, the depletion of *mHTT* in neurons, astrocytes and oligodendrocytes, excluding microglia, partially rescued the HD phenotype in BACHD mice. This demonstrates that oligodendroglia and astroglia may play an important role in the pathology of HD at an early stage of disease.

In light of this accumulating evidence that glia are affected in HD, we hypothesised that glial transcriptome signatures would be different in HD. As glia are heterogeneous across different regions of the CNS^28,29,30^, we also hypothesise that these differences might vary with region. We investigated this using snRNAseq from the post-mortem brains of people with HD and matched controls, and found abundance and gene expression changes in all glial types in the human HD brain compared to controls, in the caudate nucleus but also more globally in the cerebellum, frontal cortex and hippocampus. These changes suggest two possible HD disease mechanisms: firstly, we detect upregulation of chaperone-mediated protein folding as a cross-glial signature potentially acting as a compensatory mechanism. Secondly, we show an oligodendrocyte-specific upregulation of the calmodulin-dependent 3’,5’-cyclic nucleotide phosphodiesterase 1A (*PDE1A*), which could cause dysfunction to key cellular functions. This confirms the importance of glial responses in HD and provides us with possible targets for their manipulation for therapeutic benefit.

## METHODS

### Post mortem tissue

Fresh frozen brain samples from HD and non-neurological control donors were provided by prospective donor schemes with full ethical approval from the MRC Edinburgh Brain Bank (EBB; 16/ES/0084), the Netherlands Brain Bank (NBB; 2009/148), and Leiden University (LBB). For tissue from LBB, written informed consent was obtained for each donor in accordance with the Declaration of Helsinki and all material and data were handled in a coded fashion maintaining patient anonymity according to Dutch national ethical guidelines (Code for Proper Secondary Use of Human Tissue, Dutch Federation of Medical Scientific Societies). HD diagnosis was confirmed by neuropathologists (Table S1). Six patient donors were selected from LBB and NBB. From each donor, four different regions were included: Frontal cortex (FrCx), Caudate Nucleus (CN), Hippocampus (HC) and Cerebellum (CB). Control tissue matched for sex and age was sourced from all three brain banks. See Table 1 for summarised donor information.

**Table 1.** Summary of donor information. TAAD = Type A dissection of ascending aorta

| Brain bank | ID | sex | age (y) | pmd (h) | CAG length | Diagnosis | Cause of death |
| --- | --- | --- | --- | --- | --- | --- | --- |
| NBB | 1995-102 | m | 53 | 10:00 | na | Non-demented control | TAAD |
| NBB | 2016-056 | m | 68 | 05:50 | na | Non-demented control | Cancer |
| NBB | 2017-005 | f | 60 | 05:30 | na | Non-demented control | Euthanasia |
| NBB | 2017-124 | f | 55 | 07:30 | na | Non-demented control | Euthanasia |
| Leiden | E14-035 | f | 53 |  | na | Non-demented control | Heart failure |
| Leiden | E14-138 | m | 48 | 03:30 | na | Non-demented control | Myocardial infarct |
| EBB | BBN-15223 | m | 47 | 42:00:00 | na | Non-demented control | Coronary atherosclerosis |
| EBB | BBN-22618 | f | 45 | 40:00:00 | na | Control - schizophrenia | Suspension by ligature |
| NBB | 2013-076 | m | 71 | 04:25 | ?/43 | Huntington's disease | Euthanasia |
| NBB | 2017-060 | f | 57 | 06:40 | 18/42 | Huntington's disease | Euthanasia |
| NBB | 2017-081 | f | 38 | 05:40 | 17/54 | Huntington's disease | Euthanasia |
| Leiden | HD231 | f | 57 | 04:30 | 23/43 | Huntington's disease | Pneumonia |
| Leiden | HD240 | m | 62 | 04:30 | 20/44 | Huntington's disease | General decline |
| Leiden | HD241 | m | 48 | 02:00 | 21/45 | Huntington's disease | Euthanasia |

### Nuclear isolation

Depending on tissue size, 3-5 cryosections at 20 μm thickness were used for nuclear extraction for each tissue sample. White and grey matter was used for nuclear preparation. Extraction of nuclei was performed with the Nuclei PURE Prep Isolation Kit (Sigma-Aldrich, NUC201-1KT) according to manufacturer’s instructions. Nuclei were stained with Trypan blue, counted using an automated cell counter (Bio-Rad TC20), and normalised to a concentration of 1,000,000 nuclei per ml.

### 10X and NGS

We used the 10X Genomics 3’ gene expression V3 kits and reagents according to the manufacturer’s instructions. Samples were randomised to one of 7 different 10X Genomics Chromium single cell chip B for generation of Gel Bead-in-Emulsion (GEM). The quality of cDNA libraries was assessed using a Bioanalyzer (PerkinElmer LAS (UK) Ltd), to ensure a minimum library concentration of 5nM. Unique Illumina sequencing primers were added to each sample to allow demultiplexing of sequencing pools. Paired-end next generation sequencing of the cDNA libraries was carried out by Edinburgh Genomics. Indexed libraries were pooled together in four pools of 9 and one pool of 8 libraries. Each pool was sequenced on two lanes of an Illumina NovaSeq S2 flow cell to yield approximately 1750M read pairs per lane.

### Data preprocessing

Fastq files were aligned with the human reference genome GRCh38 using 10X Genomics CellRanger v3.0.2. Velocyto (v.0.17.16)^31^ was used to reintroduce unspliced mRNAs to the feature count matrix.

### Quality control

Quality control was carried out using Scater^32^. Genes expressed in ≤ 200 nuclei across the spliced and unspliced mRNAs were removed. Nuclei were filtered out based on high/low UMI counts, high/low gene counts and high percentage of mitochondrial genes (see Table S2 for thresholds). Doublets were removed using scDblFinder^33^, filtering out nuclei with a doublet score ≥ 0.94. After QC, we show that nuclei quality metrics were comparable to that seen in previously published data sets (Fig S2).

### Normalisation and linear dimensional reduction

After QC, the data were normalised using Scran^34^ separately for the four tissue regions, then merged and principal component analysis was performed.

### Batch correction

Canonical correlation analysis (CCA) using Seurat v4^35^ was used to adjust for batch effects. The top 2000 variable genes were selected for each 10X chip, and integration anchors found between each pair of chips, using these to carry out iterative pairwise integration using the first 37 PCs.

### Dimensional reduction and clustering

Seurat v4 was used for the dimensional reduction and clustering of cell populations. The 35 first principal components were selected to construct a UMAP. A K-Nearest Neighbour (KNN) graph was built as basis for community detection using a Louvain algorithm^36^. Initial clustering was performed to resolve broad cell types, as denoted by expression of canonical markers detailed in the main text, using a KNN resolution of 0.25. Any clusters where one donor contributed more than 50% of the cluster were removed. Clusters had to contain nuclei of at least 5 donors, while donors were only included if they contributed at least 2 % to the total cluster size. Finally, clusters consisting of fewer than 100 nuclei were removed.

### Annotation and subclustering

Subclustering for each lineage was carried out by extracting the relevant broad cluster(s), and applying the same clustering approach as above, selecting the most appropriate PC number, based on the elbow heuristic. The most appropriate KNN resolution for each cell type was selected after assessing the purity, cohesion and stability of clusters at a number of different resolutions. Neuronal populations were annotated with a number and regional specificity where appropriate, and vascular cells according to lineage and function. Glial populations were annotated with the aim to avoid giving an impression of order or rank, and to separate them from previous datasets as follows: Microglia with flower names, astrocytes with herb names, and oligodendroglia with tree names.

### Differential gene expression analysis

Differential gene expression analyses for the identification of cluster marker genes and genes altered in disease were performed using MAST^37^, filtering genes for those that had a minimum positive log2-fold change (log2FC) of 0.25 and were expressed by at least 25 % of cells within the cluster/group of interest and less than 60 % outwith. When comparing HD and control, additional thresholds of absolute log2FC of ≥ 0.8 and adjusted p-value of < 0.05 was applied. For oligodendroglia, data from the CB was excluded due to the very low number of cells captured. To display the top 5 cluster markers for each cluster of interest, the same thresholds were applied as above, selecting the 5 genes with the greatest log2FC in expression relative to other clusters.

### Network and gene ontology analysis

Protein-protein interaction (PPI) and gene ontology (GO) analysis was carried out in Cytoscape (version 3.8.2). We identified 72 genes that were shared markers of Astro_Thyme, OPC_Pine and Mglia_Violet, and used STRING to build a PPI network of the shared cluster marker genes. We then searched for functional enrichment terms associated with the genes in this network, filtering the results to include only terms related to biological processes.

### Differential abundance

Differential abundance (DA) analysis was carried out using MiloR^38^. A KNN graph was constructed and was sampled for downstream analysis using a 0.05-0.1 proportion (depending on cell type) of randomly sampled cells. The number of cells from each neighbourhood was used as input for a negative binomial generalized linear model (GML). In order to perform spatial FDR correction, the distances between each of the nearest neighbours were calculated. Comparison between HD and control was carried out with tissue region as a covariate.

## RESULTS

### snRNAseq of four regions of PM brain from HD cases and controls

We were interested in investigating the glial signatures associated with HD both in areas of MSN loss and beyond, so we carried out snRNAseq of four regions of PM brain: caudate nucleus (CN) cerebellum (CB), frontal cortex (FrCx), and hippocampus (HC). Our sample included tissue from 6 HD cases, and 7 age and sex matched healthy controls (Table 1 and S1). Following quality control of cDNA libraries, nuclei, samples and clusters, our dataset included 127,205 nuclei across 44 tissue samples, and was in keeping with quality and number from previous published datasets^39,40^ (Fig S2). We first interrogated the expression of canonical markers to identify all the major cell types of the brain (Fig 1A). We confirmed the presence of astrocytes (*ALDH1L1, GFAP*), microglia (*CX3CR1, ITGAM*), oligodendrocytes (*MOG, MBP*), OPCs and committed OPCs (COPs; *PDGFRA, GPR17*), endothelial cells (*VWF, CLDN5*), pericytes (*PDGFRB, NOTCH3*), peripheral immune cells (*CD2, CD8*), ependymal cells (*CFAP299, SOX9*), excitatory neurons (*SLC17A7*), inhibitory neurons (*GAD2*) and cerebellar granule cells (*RELN, SLC17A7*). Within the inhibitory neuron population, we identified three clusters in the CN expressing the medium spiny neuron marker *PPP1R1B*, and we confirm that the number of cells in these clusters were reduced in HD (Fig 1B) reflecting the hallmark pathology of the disease.

**Figure 1.**
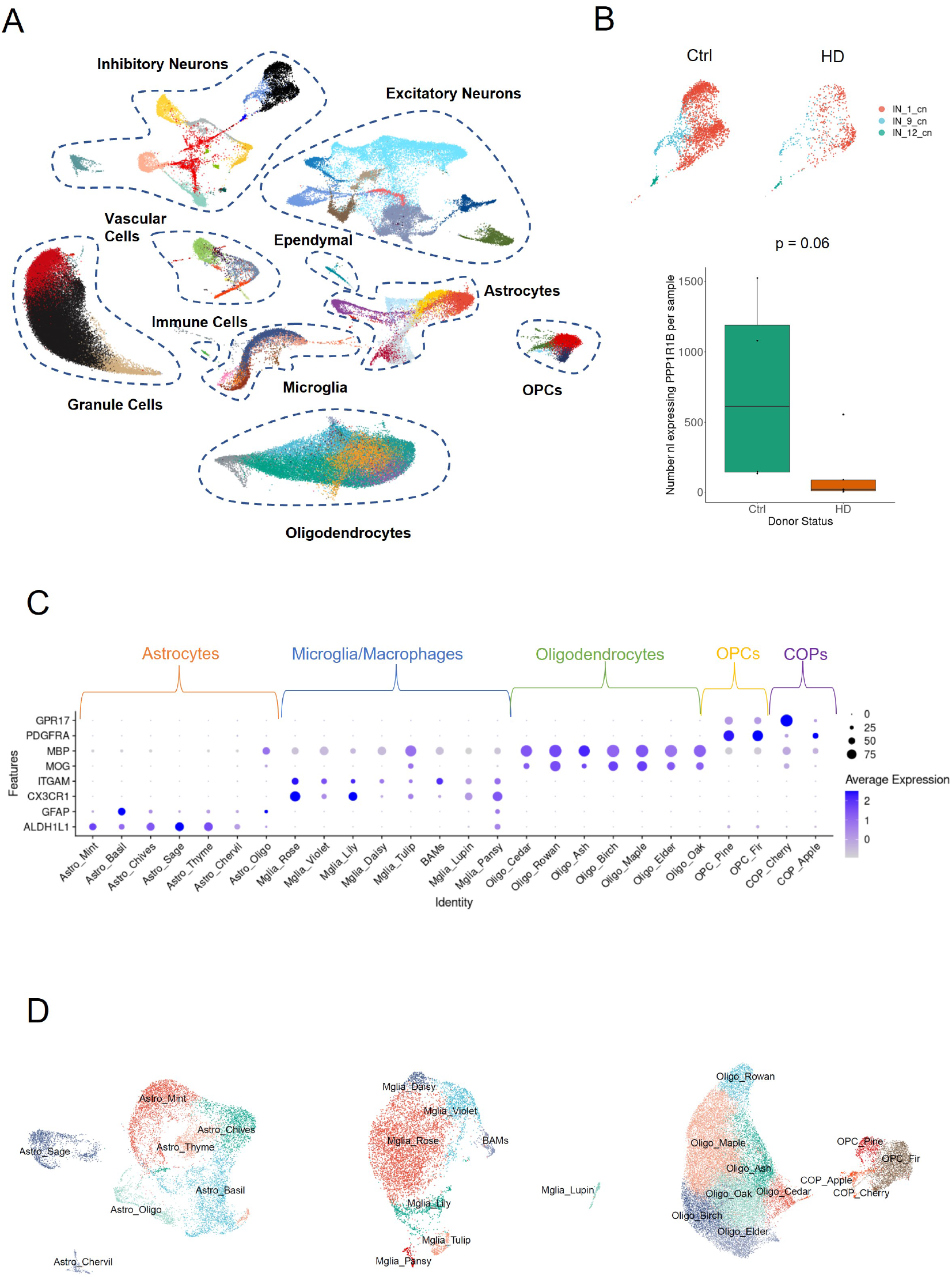
snRNAseq of four regions of PM brain from HD cases and controls. **A**. UMAP plot of all nuclei across brain regions, disease and control, showing all major cell types in the brain. **B**. Medium spiny neurons in the CN (PPP1R1B+) are depleted in HD (median nl per sample: 19) compared to control (median nl per sample: 612), Wilcoxon Rank Sum test, W = 18, p = 0.06. **C**. Expression of canonical markers used for subsetting, across glial subclusters. **D**. UMAP plots of astrocytes, microglia and oligodendroglia (L-R), showing annotated subclusters.

As the role of glia in HD pathology is less well understood than that of neurons, we here focus on the microglia, astrocytes, and oligodendroglia to provide the first comprehensive characterisation of glial changes in HD. Furthermore, the number of differentially expressed (DE) genes comparing HD and control was higher in glia than in neurons (313 DE genes across neuronal subclusters and regions, 1278 across glia). We subsetted our data for the clusters expressing high levels of canonical glia markers: microglia (*CX3CR1, ITGAM*), astrocytes (*ALDH1L1, GFAP*), and oligodendroglia (oligodendrocytes: *MOG, MBP*, OPCs: *PDGFRA*, COPs: *GPR17*), and reclustered the nuclei within these subsets. We identified 11 clusters of oligodendroglia (7 oligodendrocytes, 2 OPCs, 2 COPs), 8 subclusters of microglia, and 7 subclusters of astrocytes (Fig 1C-D). We named these clusters after plants with an aim to avoid the implication of order or rank; Microglia with flower names, astrocytes with herb names, and oligodendroglia with tree names.

### Altered glial abundance in HD

To investigate changes in the relative proportions of the identified glial subclusters between HD and control donors, we carried out differential abundance analysis across the four brain regions combined. The results showed changes in the abundance of several glial subpopulations, with some clusters enriched and others depleted (Fig 2). Most prominently, we observed a depletion of Oligo Birch (marked by *FMN1, ROR1, ITM2A, OPALIN*, Fig 2A), Mglia Rose (marked by *CX3CR1, P2RY12, TMEM163*), and Astro Oligo (marked by *PLP1, 1LRAPL1, MBP, ST18, SLC24A2*, Fig 2C), and an enrichment of OPC_Pine (marked by *VCAN, SNTG1, NRXN1, MAP2, ATRNL1*, Fig 2A), Astro_Thyme (marked by *HSPH1, SH3BGR, ATP2C1, HSPA4L, BAG3*,Fig 2C) and Mglia_Violet (marked by *HSPH1, HSPA6, DNAJB1, BAG3, CHORDC1*, Fig 2B). We assessed whether these compositional changes were driven by tissue region or uniform across regions (Fig S4). Depletion of Oligo Birch in HD was most prominent in the CN. The enrichment of Astro_Thyme in HD was most prominent in the FrCx and HD, OPC_Pine in the CB and HC, and Mglia_Violet was most strongly enriched in HC and CN.

**Figure 2.**
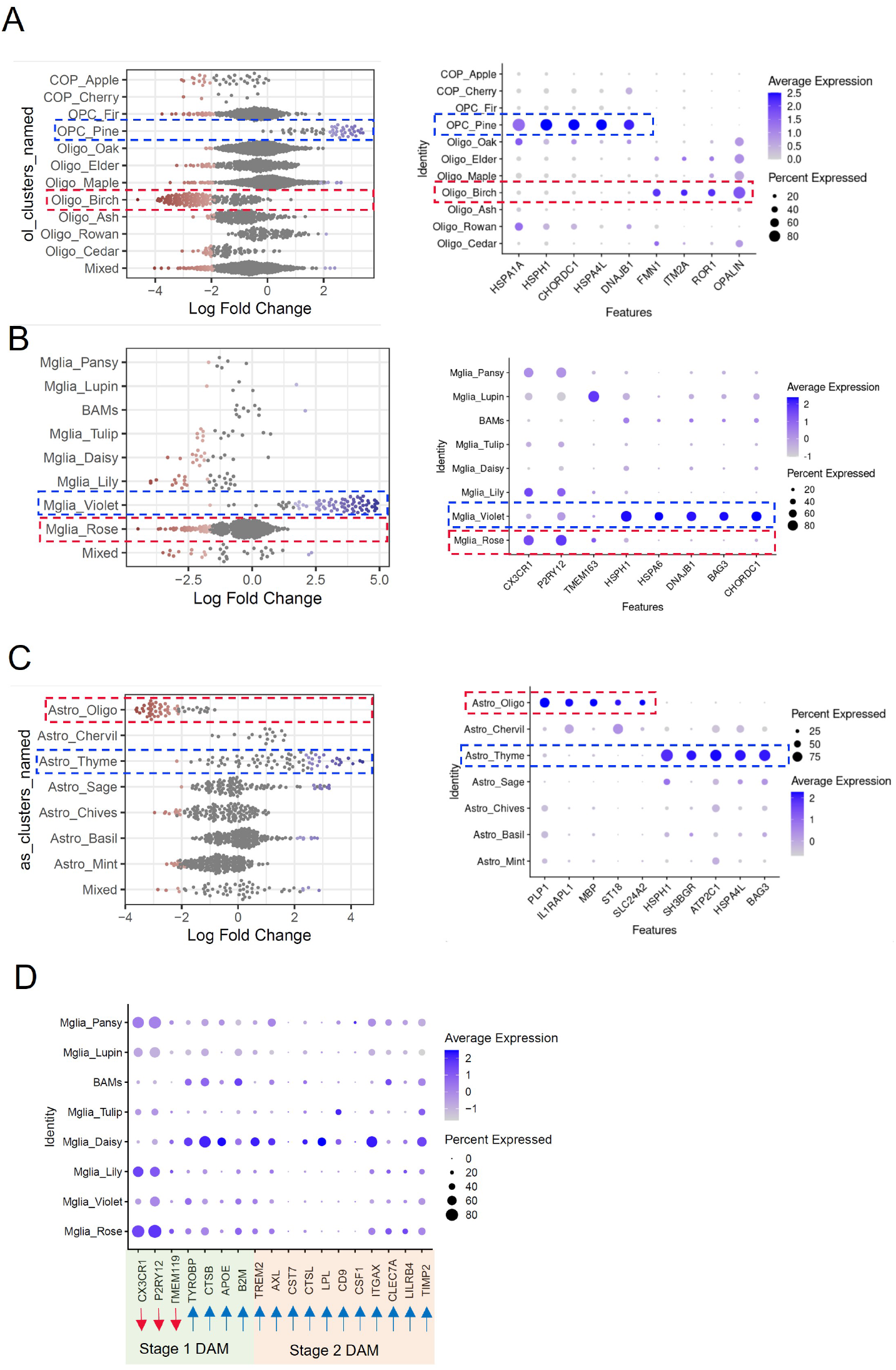
Altered glial abundance in HD. **A-C**. Beeswarm plot showing DA of subclusters of oligodendroglia (A), microglia (B), astrocytes (C), with markers of clusters that were significantly enriched (highlighted in blue) or depleted (highlighted in red) in HD compared to controls. **D**. Dot plot showing expression of stage 1 and 2 DAM markers across microglial subclusters. Red arrows denote genes downregulated in mouse DAMs, and blue arrows denote genes upregulated in mouse DAMs.

We investigated the severely depleted oligodendrocyte cluster Oligo Birch, and found that this was marked by *OPALIN*, previously described as a marker of newly formed oligodendrocytes^41^ and myelin-forming oligodendrocytes in rodents^42^, as well as the cytoskeletal component *FMN1*. The expression of these two genes in this cluster suggests Oligo Birch as an actively myelinating population of oligodendrocytes, and the loss of this cluster of myelin forming oligodendrocytes is consistent with the myelin changes seen in HD^8,20,22^. Another prominent change was the depletion of Mglia Rose, marked by *P2YRY12, CX3CR1, TMEM163*, identifying this as a more homeostatic population of microglia. As microglial activation is known to play a role in HD^7^, the depletion of this population might sugggest that a subset of microglia transition from a homeostatic to an activated phenotype.

As a disease-associated microglia (DAM) state has been described in rodent models of neurodegenerative disease and ageing^43^, we investigated whether our population Mglia_Violet, enriched in HD, was similar to the previously described DAMs by assessing the gene expression associated with stage 1 and stage 2 DAM across all microglia clusters (Fig 2D). We probed the expression of *CX3CR1, P2RY12, TMEM119*, downregulated in stage 1 DAM, and *TYROBP, CTSB, CTSD, APOE, B2M, FTH*, upregulated in stage 1 DAM. We found that the microglial cluster most closely recapitulating this pattern was Mglia Daisy, expressing low levels of *CX3CR1, P2RY12, TMEM119* and high levels of *TYROBP, CTSB, APOE* compared to other clusters. We also interrogated the expression of *TREM2, AXL, CST7, CTSL, LPL, CD9, CSF1, CCL6, IGTAX, CLEC7A, LILRB4, TIMP2*, all upregulated in stage 2 DAM. Again, the population most closely resembling this signature was Mglia Daisy, which expressed higher levels of *TREM2, AXL, CTSL, LPL, ITGAX, TIMP2* than the other clusters. However, a number of genes upregulated as part of the DAM signature such as *CD9, B2M, CSF1, CLEC7A* were more highly expressed in other clusters, and so there was no clear signature in HD microglia corresponding to the previous mouse DAM findings. Rather, our results show that the disease-associated population in this HD dataset is Mglia_Violet, which is profoundly characterised by a profile of upregulated molecular chaperone expression.

In sum, we found changes in the abundance of different glial subtypes globally across different regions in HD brain, not limited to the CN where the MSNs are lost, and with some glial clusters increased and others decreased compared to controls.

### Glia enriched in HD upregulate their expression of molecular chaperone genes

In order to better understand the differences in these glial subpopulations enriched in HD, we interrogated the marker gene expression in OPC_Pine, Astro_Thyme, and Mglia_Violet further. As the most highly upregulated genes in these three clusters appeared similar, we first probed the overlap of marker genes and found 72 genes that were flagged as marker genes for all three clusters (Fig 3 A). We then carried out functional enrichment of these genes, and found that the most significant gene ontology terms were related to protein folding (Fig3 B; Full enrichment results in Table S6). We found 10 common cluster markers related to the term “chaperone-mediated protein folding”, which included members and co-factors of the HSP70 family such as *HSPA1A, HSPA1B, HSPA9* and *ST13*, as well as those associated with the HSP90 family such as *CHORDC1* and *PTGES3*, the HSP20 family (*HSPB1*), HSP40-family (*DNAJB1*) and HSP110 family (*HSPH1*), as well as *PPID*.

**Figure 3.**
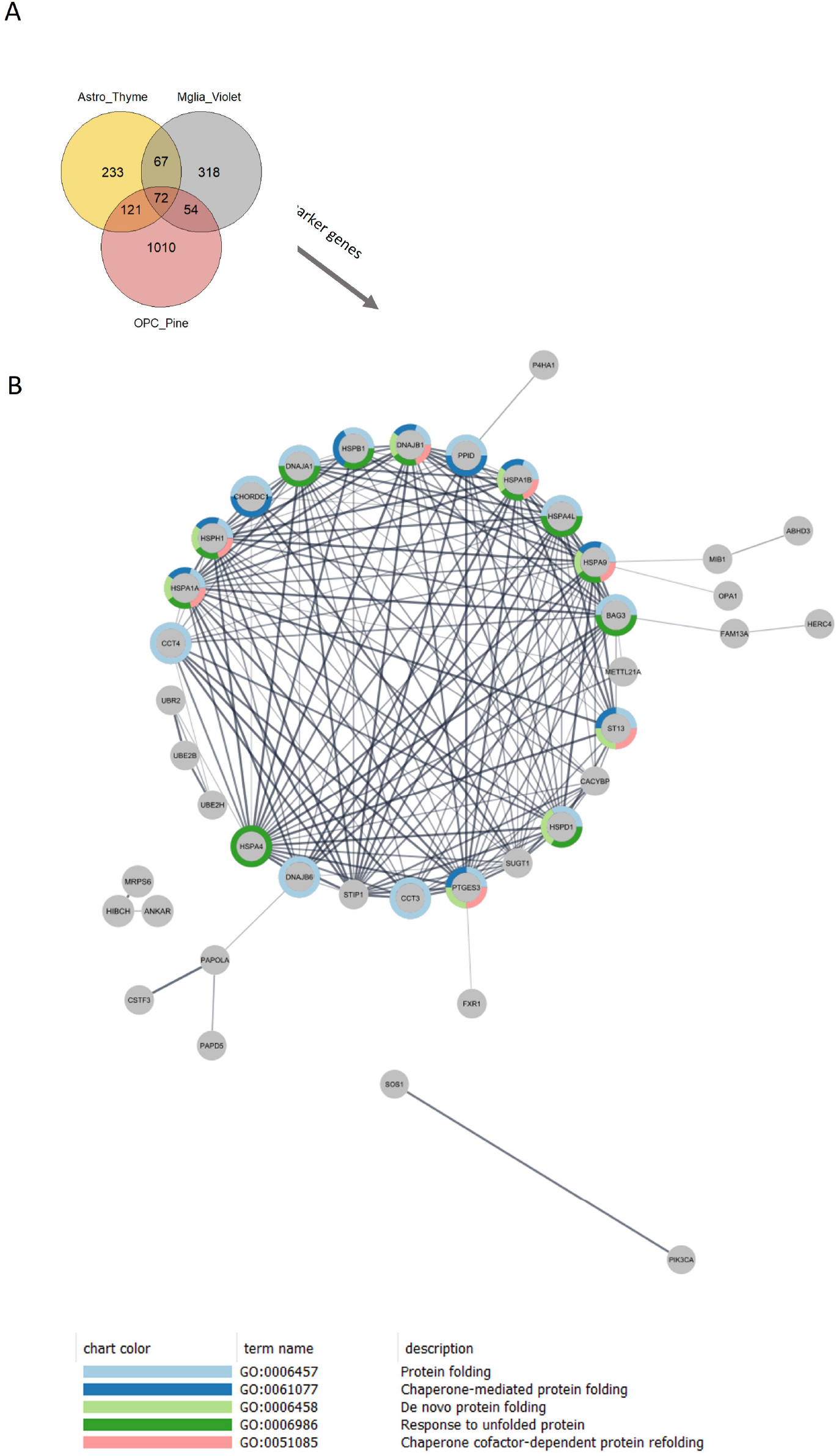
Shared cluster markers of populations upregulated in HD. A. Venn diagram showing the overlap of cluster markers between Astro_Thyme, Mglia_Violet and OPC_Pine. B. PPI network of shared cluster markers, coloured to denote those genes that belong to one of the top 5 GO terms shown below.

**Figure 4.**
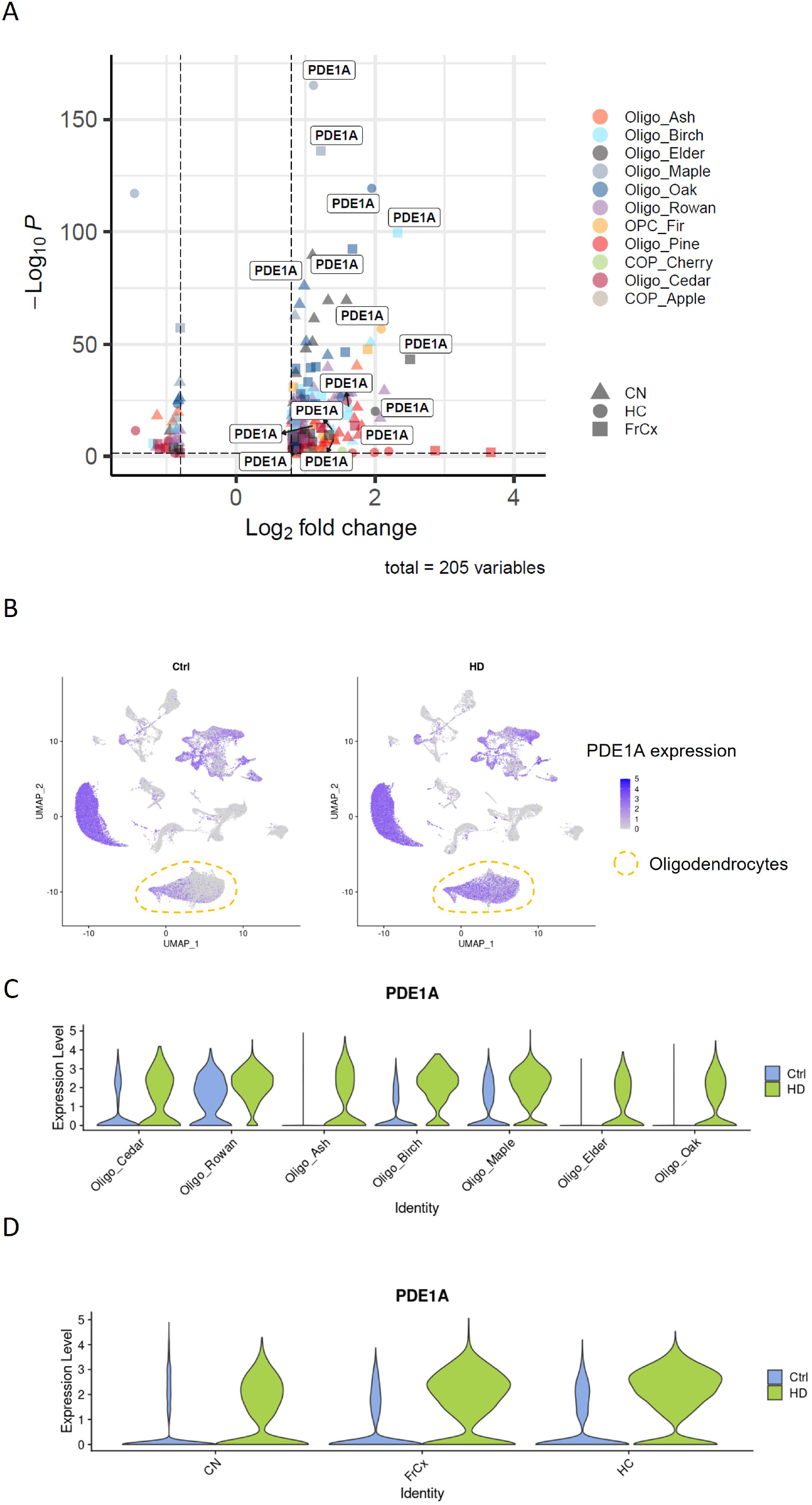
*PDE1A* upregulation in oligodendrocytes in HD. **A**. We identified upregulation of *PDE1A* across multiple clusters and regions. **B**. The upregulation of *PDE1A* in HD is only detected in oligodendrocytes. **C**. We confirmed that *PDE1A* is upregulated in all oligodendrocyte clusters except Oligo Rowan, and Oligo Ash. **D**. We observe this across CN, HC and FrCx, see main text for details.

These common gene patterns suggested a possible shared mechanism across the three glial subtypes and to investigate further, we carried out differential gene expression (DGE) analysis comparing HD and controls for each cluster of interest, applying more stringent thresholds (absolute log2FC > 0.8, FDR-corrected p < .05) for the differential expression of genes, and focusing on the brain regions where changes in abundance were most prominent (Fig S1 and S4). Considering OPC_Pine, Astro_Thyme, and Mglia_Violet separately, we observed an upregulation of molecular chaperone gene expression in HD. For Mglia_Violet, the enrichment was most prominent in the CN and the HC, where we found 136 and 440 differentially expressed genes, respectively (Table S3). Among the most highly upregulated genes in both regions were the HSP70 family member *HSPA6* and co-chaperone *BAG3*, as well as the HSP90 co-chaperone *PTGES3*. The enrichment of Astro_Thyme was most prominent in the FrCx, and here we found 14 DE genes, all of which were upregulated in HD, and the majority of these were related to chaperone-mediated protein folding (*HSP90AA1, HSPH1, HSPA1A, CRYAB, PTGES3, CHORDC1, STIP1, DNAJB1*). When looking at Astro_Thyme in the HC we found 21 DE genes (all upregulated; Table S4), including the molecular chaperone genes *HSPA4L, HSPH1, CHORDC1* and *HSPD1*. For OPC_Pine we found three upregulated genes in the HC and two in the FrCx (Table S5), including the HSP70 family members *HSPH1* and *HSPA1A*, respectively. Thus, both combined and separately in the more abundant clusters, and over different regions, we confirm upregulation of genes involved in protein folding and chaperone functions in HD vs control.This is of interest as molecular chaperones aid protein folding and in the context of stress/disease/injury, serve to prevent protein aggregation^44^ and promote degradation of misfolded proteins by the proteasome^45^. As aggregation of mutant HTT is implicated in HD pathology, the upregulation of chaperone-mediated protein folding machinery likely reflects an adaptive response, which may even promote cell survival. An upregulation of molecular chaperones has previously been reported in HD astrocytes in the cingulate cortex^46^, and we probed this further in this published dataset by investigating the expression of the molecular chaperone cluster markers shared between OPC_Pine, Astro_Thyme, and Mglia_Violet as detailed above. We found that several of these genes were expressed more highly in the clusters of astrocytes (*HSPA1A, HSPA1B, ST13, PTGES3, HSPB1*) and microglia (*HSPA1A, HSPA1B, CHORDC1, HSPB1*) that are more prevalent in HD cingulate cortex (Fig S3). To summarise, we find that upregulation of a number of molecular chaperones is a robust cross-glial signature enriched in HD, and we propose that upregulation of chaperone-mediated protein folding is a potentially beneficial compensatory disease mechanism shared across a subset of OPCs, microglia and astrocytes in HD.

### *PDE1A* upregulation in oligodendrocytes in HD

Another prominent feature in our results from the DGE analysis of glial clusters comparing HD and control was the selective upregulation of the gene encoding cyclic nucleotide phosphodiesterase *PDE1A* in oligodendrocytes. We carried out differential gene expression analysis comparing HD and control for each cluster of oligodendrocytes both with all tissues combined, and separately for each brain region. Across forebrain regions combined, *PDE1A* expression was upregulated above threshold in all clusters of oligodendroglia except Oligo Rowan and Oligo Ash, and most strongly in Oligo Birch (log2FC = 1.79, p < .001). When separating the three forebrain regions, *PDE1A* was upregulated in the clusters Oligo Oak, Oligo Elder and Oligo Birch across all regions, and in Oligo Maple, Oligo Ash and Oligo Cedar only in the HC and FrCx. Oligo Rowan (marked by *RBFOX1*) was the only cluster not showing any upregulation of *PDE1A* expression in any region, at least in part due to higher expression in this cluster in controls.While *PDE1A* was expressed in other cell types both in HD and control, we only observed changes in its expression in HD in oligodendrocytes, suggesting a pathological mechanism that is particularly relevant for this cell type, and in all brain regions examined. *PDE1A* belongs to large group of cyclic nucleotide phosphodiesterase families that hydrolyze cGMP and/or cAMP to their respective monophosphates. The PDE1 family is unique in that its activation is calmodulin-dependent, and that it has dual affinity for both cAMP and cGMP, with a higher affinity for cGMP in the case of *PDE1A*. Therefore, the upregulation of *PDE1A* in HD oligodendrocytes may limit the availability of cGMP and/or cAMP, with a potential dysregulation of downstream processes contributing to oligodendrocyte pathology in HD.

To date, only one other dataset of human oligodendrocytes in HD exists^47^, from striatal grey matter. We probed this published list of differentially expressed genes to investigate any similarities, and confirmed that *PDE1A* expression was also upregulated in oligodendrocytes in this dataset. However, the upregulation was more modest compared to our own observations, perhaps due to compositional differences between this dataset and ours.

## DISCUSSION

The present study represents the first comprehensive characterisation of glial changes across different regions of the brain in HD. The results support our hypothesis that the transcriptomic profiles of glia are altered in HD across brain regions, not limited to one type of glial cell but across microglia, astroglia, and oligodendroglia. This was true both in terms of the relative abundance of certain glial subtypes, and their gene expression profiles. The changes in glia in HD are not limited to the striatum, where there is prominent loss of MSNs, but present in all of the areas we studied with no clear regional preponderance, confirming that HD has global brain effects. As our HD donors chose euthanasia, these changes may represent earlier brain pathology than other studies using post mortem brain of end stage disease patients. We highlight two prominent molecular pathways upregulated in a subset of glia in HD, which provide insight into pathological mechanisms and novel therapeutic targets: chaperone-mediated protein folding and phosphodiesterase activity.

The importance of molecular chaperones in neurodegeneration has been explored previously^45^ as they play an important role in maintaining proteostasis both in healthy cells and in the context of disease. Some HSPs and co-chaperones are constitutively expressed, but others are inducible by stress/disease/injury and have been shown to be altered in neurodegeneration. The increased presence of these in our dataset is not likely to be simply technical, through cellular stress, since these nuclei pass quality control, it is selective for some cellular clusters and not others, and is not found in the control donors undergoing the same processing. In mouse, fly and yeast models, overexpression of HSP70 has been shown to suppress polyQ-aggregation and subsequent neurodegeneration^48,45^, and the balance between HSP90/HSP70 may be a key mechanism - supported by the finding that upregulation of HSP70 and inhibition of HSP90 both reduce protein aggregation and toxicity^45^. Furthermore, HSP-genes have been previously linked with HD^46^, vascular dementia^49^ and Parkinson’s disease^50^, reflecting its relevance to neurodegeneration. The upregulation of molecular chaperone genes in HD is likely an adaptive response to the aggregation of mutant HTT, and the higher relative abundance of cells expressing this machinery suggests it may promote cell survival.

Conversely, the upregulation of *PDE1A* expression in oligodendrocytes in HD likely represents a detrimental mechanism as this inhibits important downstream processes through its regulation of cAMP/cGMP. The inhibition of various phosphodiesterase family members has previously been investigated in the context of both oligodendroglial functioning and HD. Inhibition of PDE7^51^, PDE4^52^ and PDE3^53^ in rodents *in vivo* and/or *in vitro*, increased oligodendroglial differentiation and/or survival, as well as remyelination and rodent behavioural performance after demyelination. These studies in development, injury and disease highlight the potential for phosphodiesterases as a therapeutic target to promote remyelination in neurological conditions. Inhibition of a different phosphodiesterase, PDE10A, in animal models of HD improved striatal pathology and behavioural performance^54^, and restored the functioning of cortico-basal ganglia pathways supporting motor performance^55^. However, a randomised double-blind clinical trial of a PDE10A inhibitor in people with HD failed to improve motor performance^56^. *PDE1A* has mostly been studied in the heart, where it exerts its role primarily through regulation of cGMP levels^57^, and where inhibition of *PDE1* may be of therapeutic benefit in heart failure. In a mouse model of cardiac proteinopathy, *PDE1A* was upregulated and PDE1 inhibition promoted proteasomal degradation of a misfolded protein, improving cardiac function and survival^58^.

The transcriptomic heterogeneity of glia has also been demonstrated in other neurological disorders, including multiple sclerosis^39^, Alzheimer’s^40^, Parkinson’s^59^, and major depressive disorder^60^, and the question remains whether the changes in the abundance of different clusters represent the selective death of some subtypes of glia and the survival of others, or whether it represents a dynamic ability of the cells to change their transcriptomic “state” in response to disease or injury. Our finding of a cross-glial signature of upregulated chaperone-mediated protein folding provides some evidence of the latter as this likely represents an adaptive response to the presence of *mHTT* in HD. The lack of a clear rodent DAM signature microglial cluster in HD is consistent with findings from a single-cell RNAseq study of Alzheimer’s patients^61^, where there was also no single microglial cluster expressing the rodent DAM signature, and a more disperse expression of DAM signature genes across different microglial clusters.

These findings of global changes in glia in HD support a therapeutic disease-modifying strategy with growing support in several neurodegenerative diseases that targeting a pathological mechanism within a glial population may be beneficial as this is more tractable than targeting neurons. Therefore, our study highlights an opportunity for novel therapeutics in HD.

## Supporting information

Boestrand.et.al_supplementary_tables

## ACKNOWLEDGEMENTS

We thank the MRC Edinburgh Brain Bank, Netherlands Brain Bank and Leiden University for providing post mortem tissue. We thank the following people and facilities at the University of Edinburgh for their help: Pamela Brown from the Biomolecular Core facility for Bioanalyzer measurements, Andrea Corsinotti for providing materials for cDNA library preparation. Next Generation Sequencing of cDNA was carried out by Edinburgh Genomics at the University of Edinburgh. This work has made use of the resources provided by the Edinburgh Compute and Data Facility (ECDF) (http://www.ecdf.ed.ac.uk/). We also thank neuropathologists S.G. van Duinen at the University of Leiden and Colin Smith at the University of Edinburgh.

## CONTRIBUTIONS

SMKB conducted the snRNAseq experiment with assistance from LAS, SJ, and NLK. SMKB analysed the data with support from LAS, and input from NBC, NLK, and AW. NCH provided access to a 10X chromium controller and reagents. WvRM and SdB, with help from BK, provided access to post mortem tissue from Leiden University. AW, JP and LAS supervised the project. SMKB and AW wrote the first manuscript version and all authors contributed with their comments to the submitted version.

## SUPPLEMENTARY FIGURES

**Figure S1.**
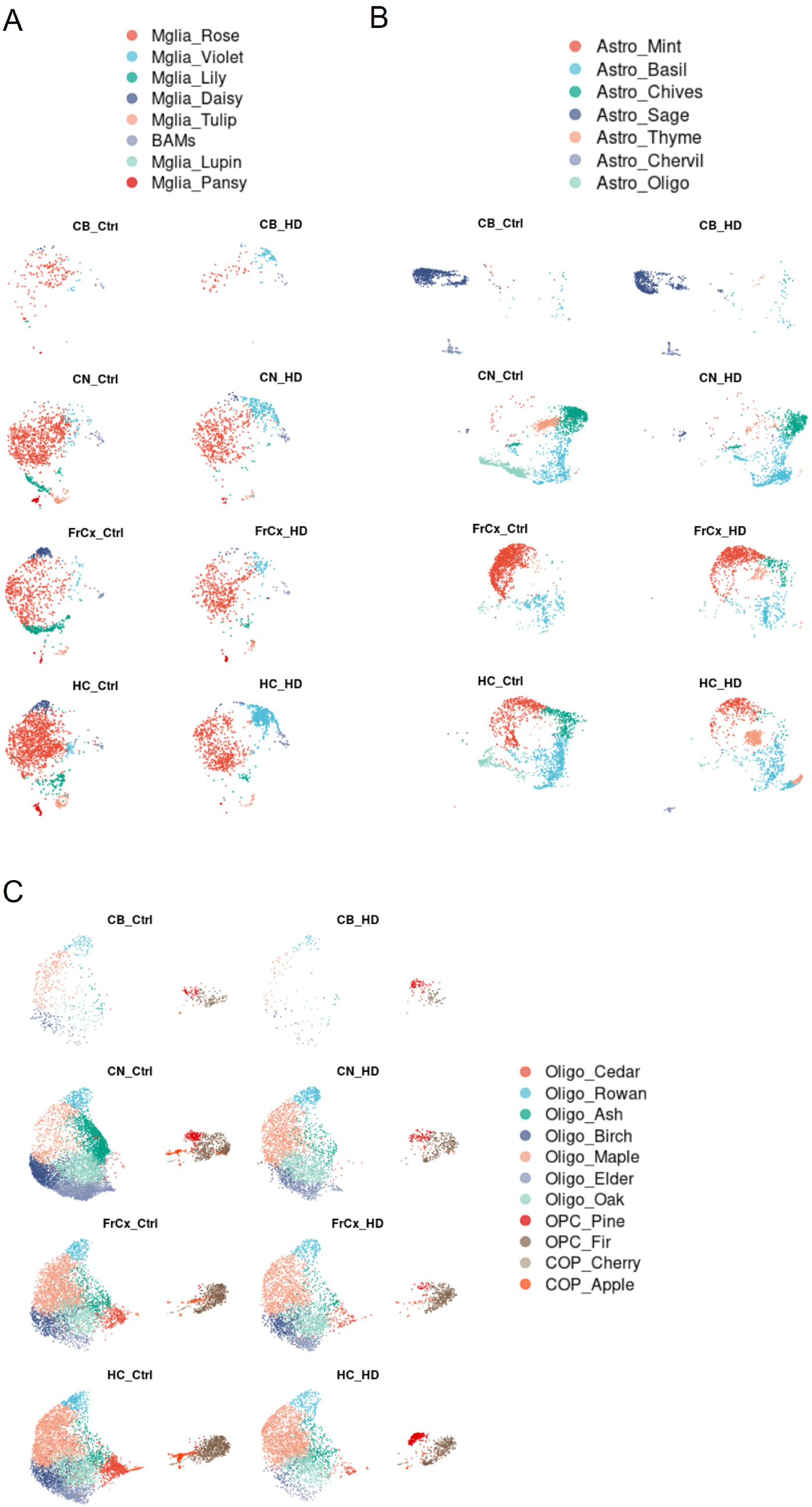
UMAP of glial subclusters, separated by region and condition. A. Microglia, B. Astrocytes, C. Oligodendroglia

**Figure S2.**
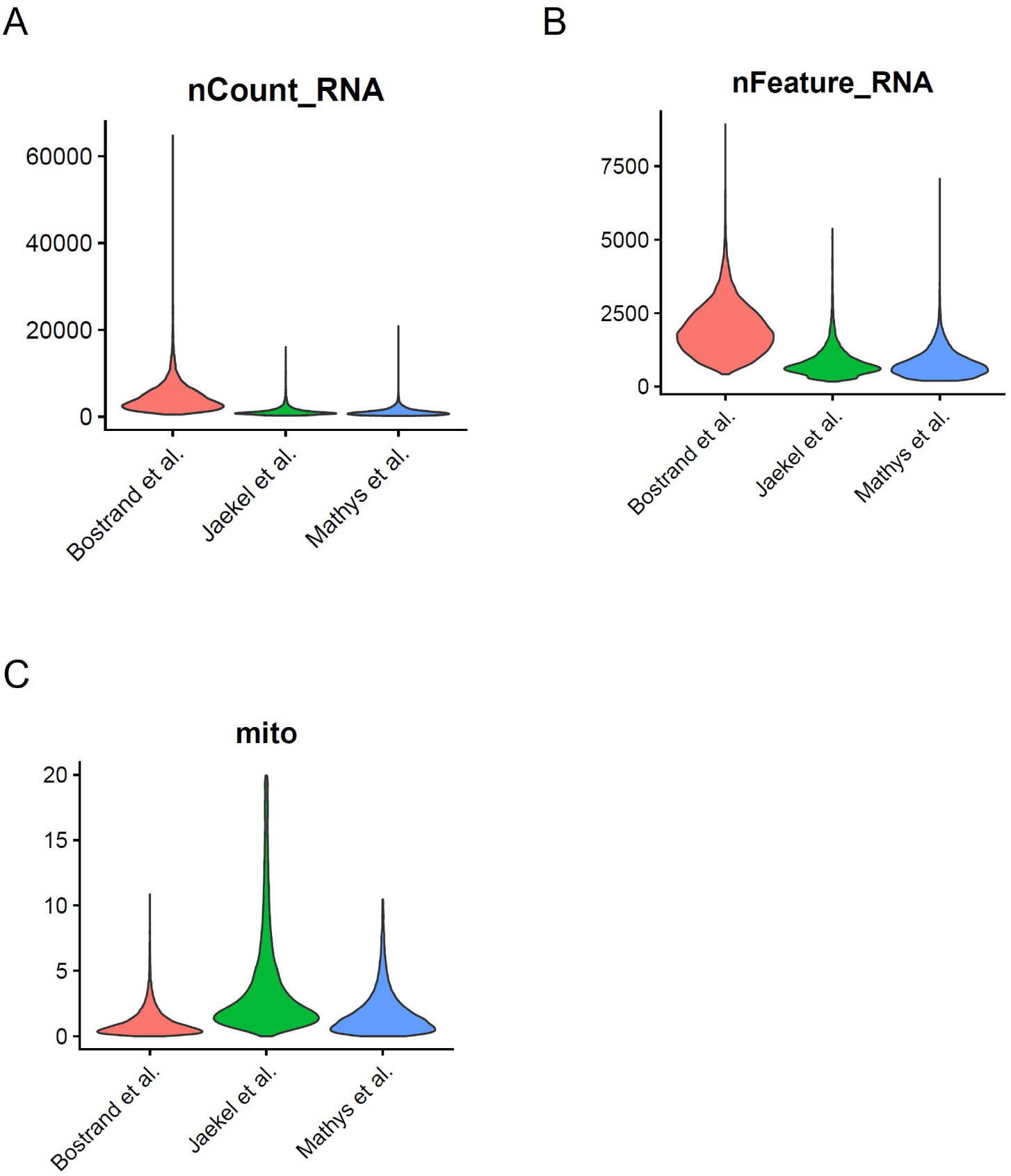
Comparison of nuclei quality with two published datasets. A. UMI count, B. Gene count, C. Percentage of mitochondrial genes

**Figure S3.**
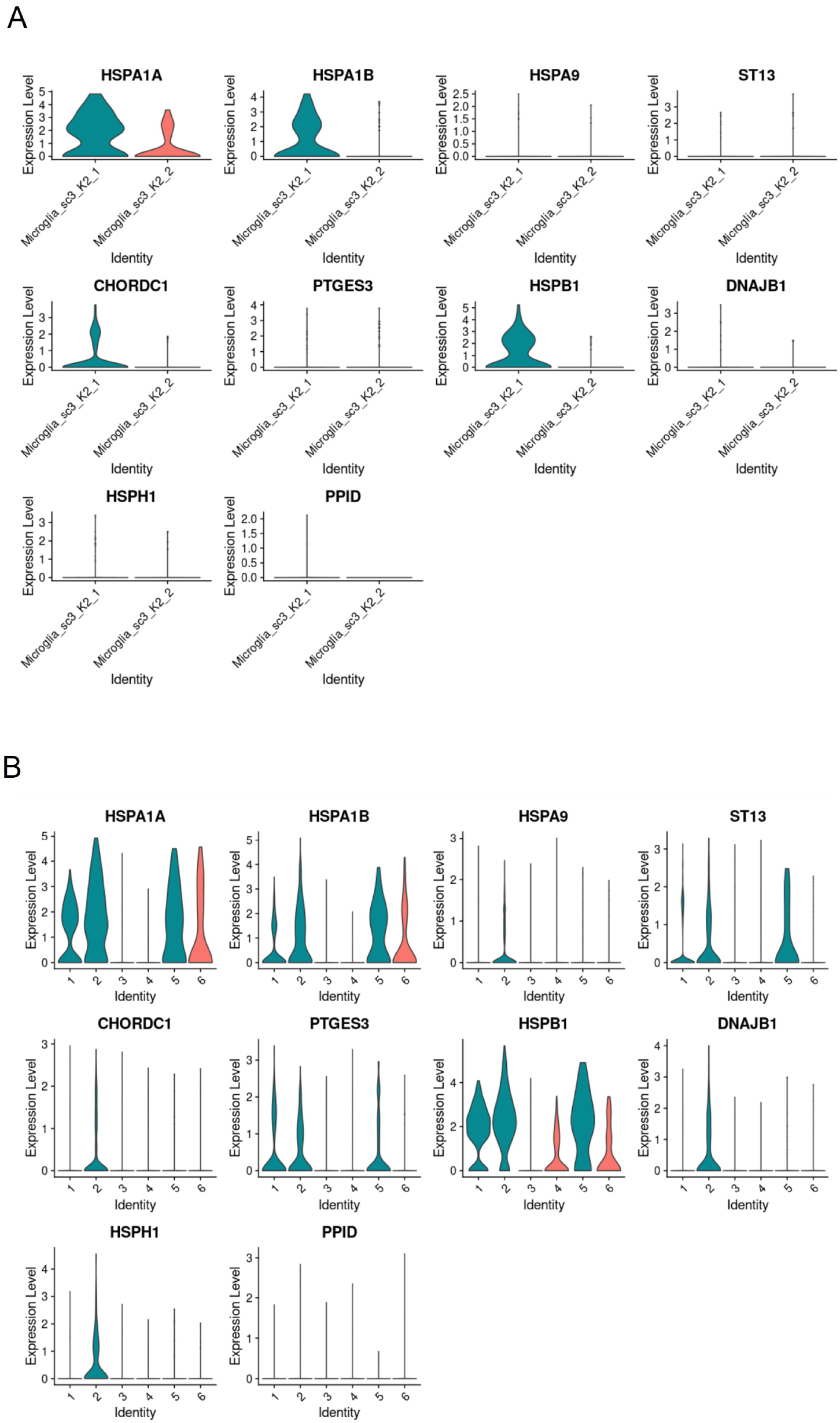
Molecular chaperones detected as upregulated in Astro_Thyme, OPC_Pine and Mglia_Violet in our dataset are also upregulated in microglia and astrocytes from Al-Dalahmah et al.^46^. A. Expression in microglia, where “Microglia sc3 K2 1” is enriched in HD, B. Expression in astrocytes, where populations 1,2 and 5 are enriched in HD.

**Figure S4.**
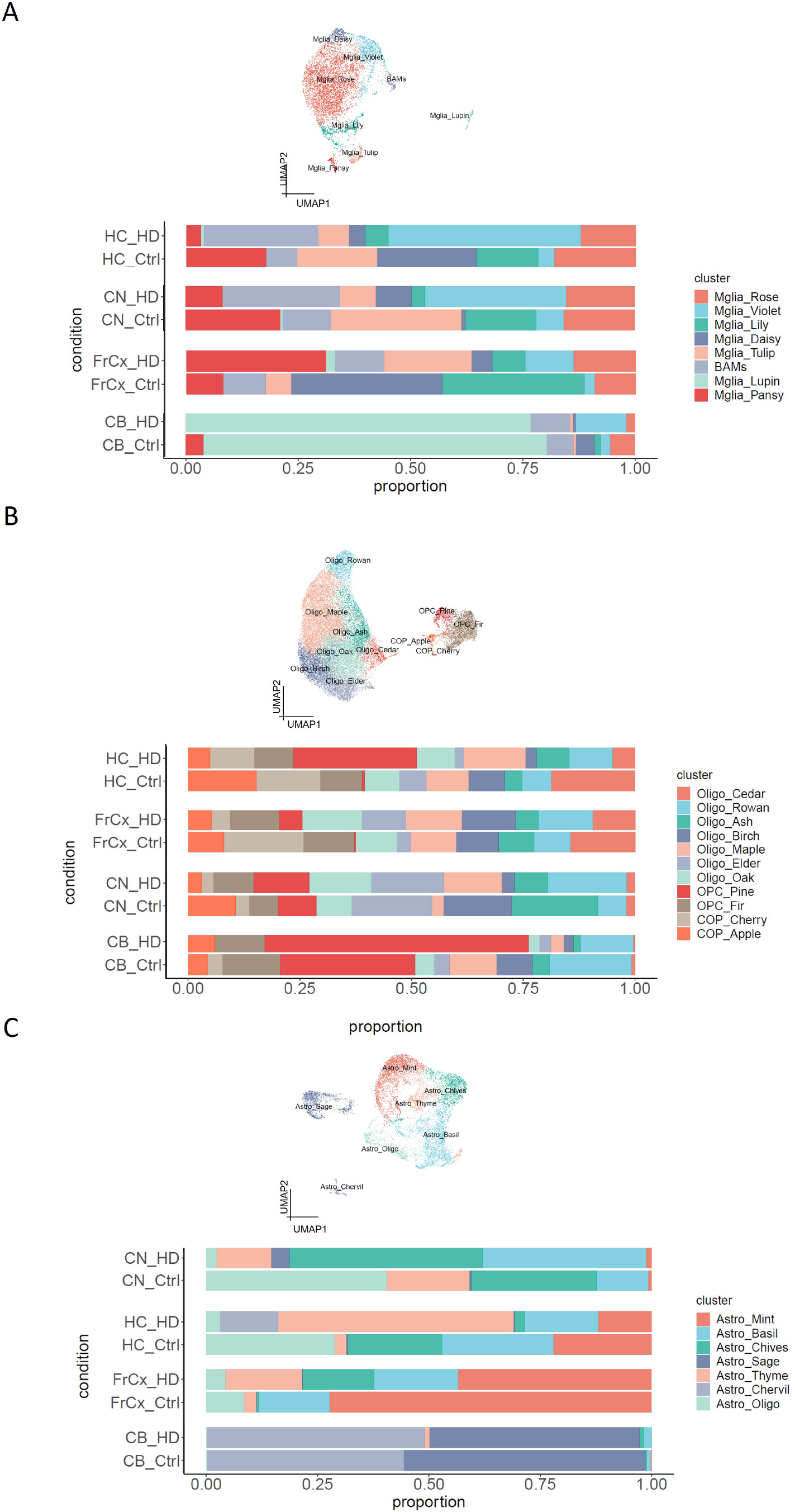
Cluster proportions comparing regions and conditions. A. Microglia B. Oligodendroglia, C. Astrocytes

